# Collateral effects of drug resistance evolution: explaining repeatability and directionality patterns

**DOI:** 10.64898/2026.03.26.714451

**Authors:** Maya Louage, Barbora Trubenová

## Abstract

Drug resistance evolution is hindering the treatment of various diseases where new drug development cannot keep pace. One possible solution to this problem could be to exploit trade-offs in resistance to already existing drugs. One such trade-off is collateral sensitivity, where resistance to one drug increases susceptibility to another. Contrarily, cross-resistance means that resistance to one drug implies resistance to another. However, the evolution of these two collateral effects is not always repeatable, nor is it always independent of drug order (i.e., bidirectional). Understanding what drives repeatability and directionality would help define the clinical applicability of collateral sensitivity and avoid cross-resistance. Nevertheless, the genetic and evolutionary mechanisms causing non-repeatability and unidirectionality patterns are not yet fully understood. In this study, we aim to define which drug concentrations, population dynamics, and genetic architectures give rise to these patterns of collateral effects. We describe minimal examples of the fewest loci and conditions needed for repeatable and non-repeatable, uni-or bidirectional cross-resistance and collateral sensitivity to occur. We show that clonal interference can explain unidirectional cross-resistance and can increase repeatability, whereas a strong selection, weak mutation regime increases non-repeatability. Additionally, we show how drug concentration can change the collateral effect and/or its pattern. Overall, we show how non-repeatability and unidirectionality of collateral effects are not properties solely of a drug pair but also of the selection regime: drug dose and population dynamics. We suggest that further studies, combining mathematical modelling with measurements of mutations and full dose-response curves for drug pairs with known patterns of collateral effects, are needed to understand this problem.

## 1 Introduction

In this article, we focus on the evolution of collateral effects, which result when adaptation to one drug alters susceptibility to another. Collateral effects can be either beneficial to the pathogen, termed cross-resistance, or detrimental to it, referred to as collateral-sensitivity (Casier et al., 2025). Cross-resistance can facilitate multidrug resistance, and it is essential to consider it in order to optimise drug treatment. Cross-resistance can occur when drug pathways overlap between two drugs, or when a mutation influences multiple pathways (pleiotropy), which confer resistance to different drugs (Alekshun and Levy, 2007). A specific example of the former is a mutation of a multidrug efflux pump regulator in *Pseudomonas aeruginosa* (Cao et al., 2004).

On the other hand, collateral sensitivity may be exploited by using drugs cyclically or sequentially to improve treatment success (Nichol et al., 2015; Aulin et al., 2021; Kim et al., 2014). However, our ability to exploit collateral sensitivity is hindered by our limited ability to predict it (Casier et al., 2025). Collateral sensitivity may result from several mechanisms, including increased drug toxicity or uptake (Roemhild et al., 2020).

Collateral effects of drug resistance evolution receive most attention in studies of bacteria and antibiotic resistance, even though these effects are general and have been documented in multiple classes of pathogens as well as in cancer (Hellinga et al., 2024; Carolus et al., 2024; Ogbunugafor et al., 2016; Zhao et al., 2016). Primarily, studies of collateral effects use large-scale repeated experimental adaptation to different drugs. A parental bacterial strain is evolved in one drug environment, and then the response of one or a few adapted colonies is measured against a different drug. These studies show diverse collateral effect patterns, particularly in their repeatability, directionality, and temporal dynamics (Sakenova et al., 2025; Maltas and Wood, 2019; Maltas et al., 2025; Ĺazár et al., 2014; Nichol et al., 2019; Imamovic and Sommer, 2013; Barbosa et al., 2017; Láźar et al., 2014). Experimental studies often identify mutations conferring resistance and quantify collateral effects as the shift in the minimum inhibitory concentration (MIC) relative to the parental strain.

Some experimental adaptation studies have investigated selection regimes by changing the gradient at which the drug concentration is increased (Brepoels et al., 2022; Chauhan et al., 2025) and additionally, also the maximal drug concentration (Oz et al., 2014; Jahn et al., 2017) or by only changing the maximal drug dose by phenotyping an adapting population at different time points of the same experiment (Maltas et al., 2025). Another study exposed *P. aeruginosa* to two constant sublethal drug concentrations (Sanz-Garćıa et al., 2020). No consistent results were found across these studies, nor were the findings explained within each study. Modelling studies that include collateral effects and compare treatment strategies rarely consider changes in selection regime (Aulin et al., 2021; Nichol et al., 2015; Hobbs et al., 2023) (for exemptions, see e.g. (Ardell and Kryazhimskiy, 2021)).

Despite the importance of collateral effects of drug resistance across clinical and agricultural settings, the evolutionary conditions combined with the genetic architecture that lead to collateral effect patterns of repeatability, directionality (sometimes denoted reciprocity), and their temporal dynamics are not yet fully described. In this study, we propose a simple theoretical framework to explain multiple experimental observations: uni-and bidirectional, as well as repeatable and non-repeatable cross-resistance and collateral sensitivity. Our model assumes that all changes in drug response are genetic and heritable, and not physiological (Roemhild et al., 2018). Our reasoning links dose-response curves (pharmacodynamics) to multi-locus population genetics. We demonstrate our reasoning using minimal examples with the fewest loci needed to explain these patterns and discuss the underlying assumptions. Moreover, we show that the temporal dynamics of these collateral effects are a natural consequence of our assumptions.

Importantly, we explore how evolving pathogen populations in different drug concentrations can lead to distinct collateral effects and/or patterns depending on their genotype-phenotype maps. This can also depend on the population and mutation dynamics, here denoted as the selection regime: strong selection, weak mutation regime (selective sweeps), and clonal interference (competition between mutants).

## 2 Model framework

We base our model on a combination of pharmacodynamics and population genetics, as reviewed in Witzany et al. (2023). This modelling approach has already been applied to a single-drug environment for models of drug resistance evolution in bacterial biofilms and worms (Trubenová et al., 2022, 2025) as well as to a two-drug environment (Engelstädter, 2014).

### Definitions of patterns of collateral effects of drug resistance

We define a collateral effect as a shift in the minimum inhibitory concentration (MIC) value for a drug other than the one to which the pathogen has adapted. This collateral effect is then classified as either cross-resistance, collateral sensitivity, or no collateral effect. For different pathogens, resistance might be quantified differently. We chose MIC as collateral effects are most studied in bacteria. For clarity and simplicity, we will refer to the two drugs as drug A and drug B in the following text, without further specification of their exact nature. The first pattern is repeatability (Nichol et al., 2019). A repeatable collateral effect occurs when evolution is repeated from the same parental strain under the same conditions in drug A, and the adapted resistant strains *consistently* show *the same* collateral effect when exposed to drug B. On the other hand, non-repeatability implies that the adapted strains display a *different* collateral effect or no collateral effect at all. The second pattern is bidirectionality (Sakenova et al., 2025). A collateral effect is bidirectional if adaptation to drug A or drug B results in the same collateral effect in the other drug, regardless of which drug is used first. Contrarily, unidirectionality means that only adaptation to drug A leads to a particular collateral effect in drug B, but the reverse is not true.

### Pharmacodynamics

is used to describe the relation between drug concentration and some fitness measure of a population, e.g., its growth rate. It is captured by a dose-response curve. For bacteria, these curves can take the following form (Equation 1 and Figure 1) (Regoes et al., 2004):

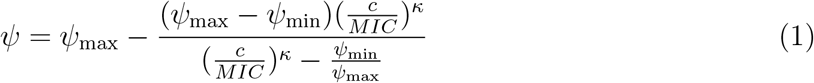

**Figure 1:**
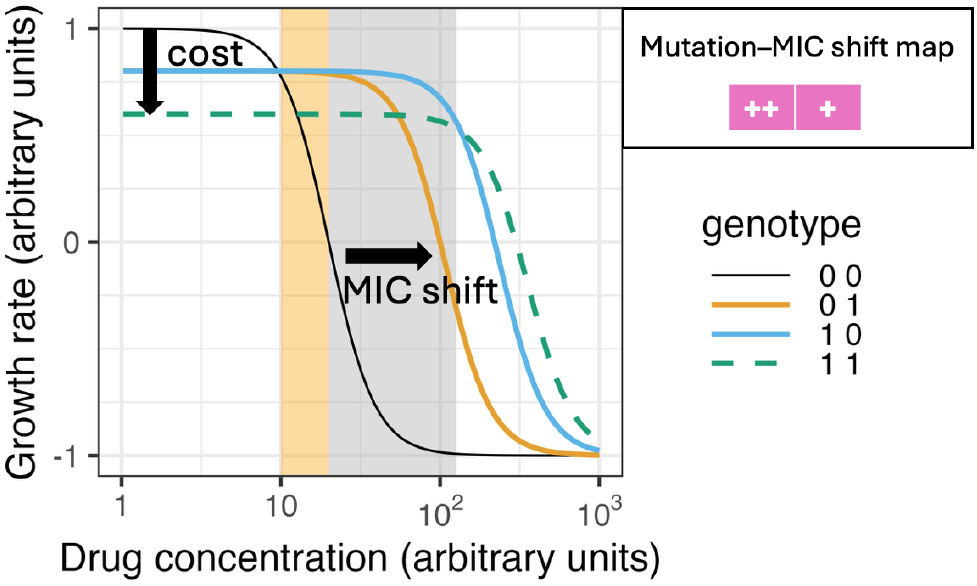
Dose-response curve example for two-locus drug resistance. Mutations that decrease MIC are not shown. Two mutant selection windows (MSW) are depicted in orange and grey. In the table (top right), each column is a locus that can be mutated. Mutational effects are shown by the symbols +, indicating resistance, and ++, indicating a larger MIC increase than +. Hence, the [0 1] genotype has a lower MIC than [1 0].

where *ψ* is the net growth rate, *ψ_max_* is the maximal growth rate in the absence of a drug, *ψ_min_* is the minimal growth rate (often negative), *c* is the drug concentration, *MIC* is the minimum drug concentration required to inhibit growth (i.e. net growth rate of the population is zero) and *κ* defines the steepness of the decline in growth rate as drug concentration increases. While often used to describe the growth of bacteria in an environment containing antibiotics, the dose-response curve can be modified for other organisms, such as insects and parasitic worms, for example, to describe the survival probability or inhibition of larval development (Caprio, 1998; Dobson et al., 1996). Our illustrations will use the bacterial dose-response curve, as most research on collateral effects is conducted in this field. The same logic presented here can be applied to other versions of dose-response curves.

### Population genetics linked to pharmacodynamics

describes the effect of mutations on the dose-response curve. We assume that resistance can be encoded by multiple loci (Nichol et al., 2019; Ogbunugafor et al., 2016), and that each locus has only one possible mutation, i.e., there are two alleles per locus: wild type, represented by 0, and mutant, represented by 1. Any mutation corresponds to a possible shift in MIC to a given drug, either as an increase relative to wild type (resistance), or a decrease (increased sensitivity). Mutations can shift the MIC in different manners for different drugs. For instance, a mutation in one locus can increase the MIC for one drug, while leaving it unchanged, decreasing, or increasing it for another. We will show this link using mutation-MIC shifts (specific genotype-phenotype maps, represented as tables with pink and green row colours, as shown in examples in Figure 1 and all scenarios in Figure 2). These depict how mutations at different loci affect the MICs of different drugs. An empty cell refers to the absence of an effect on the MIC for this drug when mutating this locus. Additionally, mutations may reduce the maximum growth rate *ψ_max_* (cost) (Andersson and Hughes, 2010), but this is not always the case (Dunai et al., 2019; Vogwill and MacLean, 2015).

**Figure 2:**
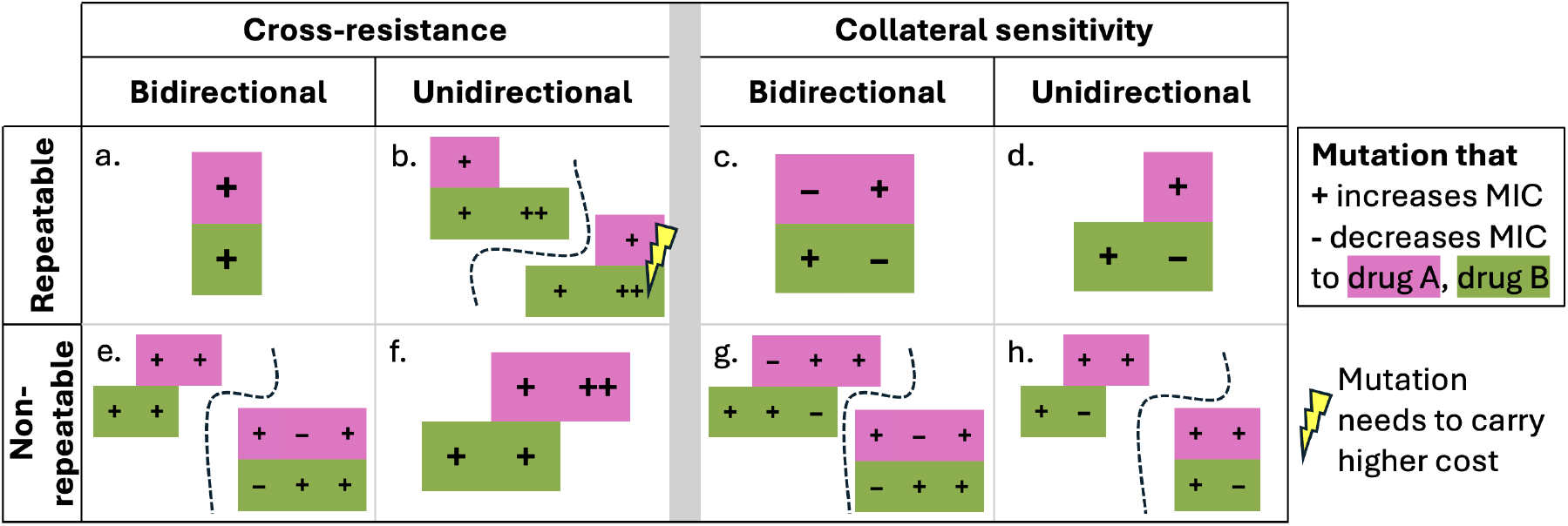
Overview table of the fewest loci with mutational effect size needed to explain directionality and repeatability of collateral sensitivity and cross-resistance depending on drug dose, cost, and selection regimes. In each cell, mutation-MIC shift tables are shown; columns represent mutable loci; rows indicate drug-specific effects of these mutations: drug A, pink top row, and drug B, green bottom row (+ means resistance, ++ means stronger resistance and - means sensitivity compared to wild type). Note that for visual clarity, the higher cost scenario in cell b was omitted in cell f. The scenario where cross-resistance is repeatable in one direction, not in the other, is not explicitly labelled but can occur with the mutation-MIC shift tables in cell b (Sections 5 and S1.1).

The dose-response curves for all genotypes define the mutant selection window (MSW). The MSW is a drug concentration range where some mutant(s) outcompete the wild type, have a positive growth rate, and are thus selected for. In Figure 1, the orange MSW shows the concentration range where both single mutants are similarly fit, and the double mutant escapes extinction. The grey MSW shows where the high-resistance single mutant (blue) is fittest. Here, also the double mutant and, at low concentrations, the low-resistance single mutant escape extinction.

In all our minimal examples, we assume that multiple mutations combine additively to change the two aforementioned parameters, *MIC* and *ψ_max_*, thus modifying the dose-response curve (parameter values are given in Table S1). For simplicity, we additionally assume that the steepness of the dose-response, *κ* (Das et al., 2020; Chevereau et al., 2015), and minimal growth rate, *ψ*_min_, are not affected by mutations. We use these assumptions in our illustrations. However, only the qualitative characteristics of the dose-response curves, e.g., mutant selection windows, matter for our reasoning. These characteristics can hold even when the aforementioned assumptions are violated (Section S1.2).

## 3 Minimal examples explaining patterns of collateral effects

Below we present minimal examples that use the fewest loci needed to explain (non-)repeatability and uni-/bidirectionality. We consider adaptation to a single drug A at a specific concentration and the collateral effect of the resulting adapted strain when exposed to drug B (A *→* B) and vice versa (B *→* A). We consider drug concentrations where a single mutant is fittest due to resistance costs. Occasionally, we discuss no-cost scenarios where the double mutant is similarly fit at certain concentrations. We assume mutation rates are high enough to prevent extinction at least sometimes. We consider collateral effects after the population’s genotype composition stabilises. Section 4 relaxes this assumption.

### 3.1 Only bidirectional, repeatable cross resistance can be explained by a single locus

The simplest scenario to explain is **bidirectional, repeatable cross-resistance**, where a single locus confers resistance to both drug A and drug B (Figure 2a, Figure S1a). Evolution in an environment containing either drug at a concentration within the mutant selection window (MSW) will lead to this mutant outcompeting the wild type and fixing in the population in any repeated experiment with successful adaptation. Therefore, this scenario describes repeatable and bidirectional cross-resistance. This, or a similar structure in which a single locus or trait correlates resistance to one drug with resistance to another, has been used in models of insecticide resistance evolution (Caprio and Suckling, 2000; Hobbs et al., 2023). The above mentioned reasoning can be extended to more loci as long as all mutations conferring resistance to drug A also confer resistance to drug B, and vice versa.

In contrast, a scenario where adaptation to either drug always confers collateral sensitivity to the other drug, denoted **bidirectional, repeatable collateral sensitivity**, requires at least two loci. A mutation at one locus gives resistance to drug A and collateral sensitivity to drug B, and at the other locus it gives resistance to drug B and collateral sensitivity to drug A (Figure 2c, Figure S1b). Selection by either drug on a wild type population will select for the mutation that gives collateral sensitivity to the other drug, as this is the only resistance mutation available. This thus leads to bidirectional, repeatable collateral sensitivity.

### 3.2 Unidirectional cross-resistance can be explained by low and high-resistance mutants

When a collateral effect is only present in the direction of evolving the pathogen in drug A and exposing it to B (A *→* B), and not drug B to A (B *→* A), it is called unidirectional.

#### Unidirectional, repeatable cross-resistance

can be explained by assuming that the mutation conferring resistance to drug A *is* associated with low resistance to drug B, while the mutation that carries high resistance to drug B *is not* associated with resistance to drug A (Figure 3a, summarised in Figure 2b). This scenario thus implies that the cross-resistance to drug B, observed when the population is exposed to drug A, is low compared to the resistance that could be achieved if the population had adapted to drug B.

**Figure 3:**
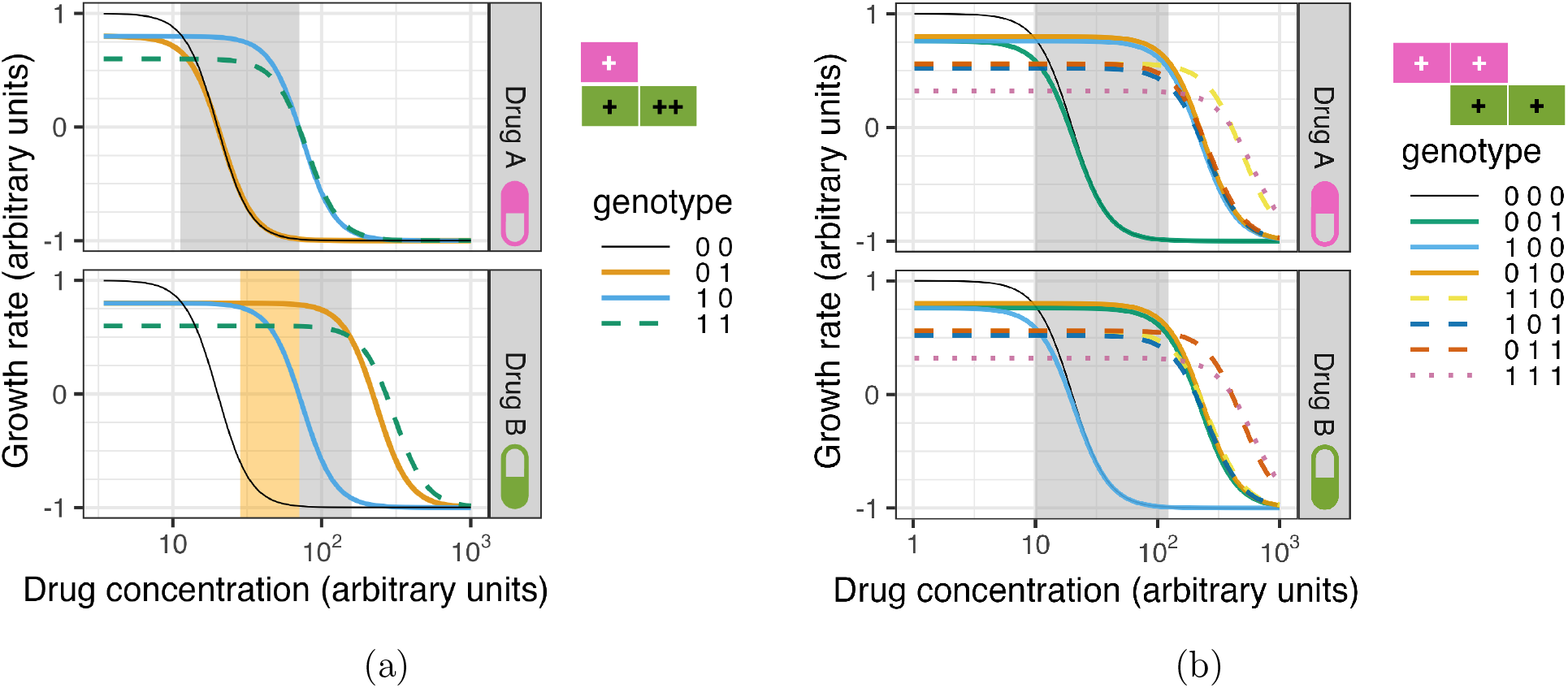
Dose-response curves for the same genotypes in two different drugs, explaining uni-directional and non-repeatable cross-resistance. In the mutation-MIC shift table (top right), columns represent mutable loci; rows indicate drug-specific mutational effects (+ means resistance, ++ means stronger resistance). **(a) Unidirectional, repeatable cross-resistance can arise when a low-resistance mutant confers cross-resistance, but a high-resistance mutant does not.** Grey shading marks concentrations where cross-resistance occurs only when adapting to drug A ([1 0] genotype), not drug B ([0 1] genotype). Orange shading for drug B shows where competition fixes the genotype without a collateral effect in drug A ([0 1] genotype). **(b) Non-repeatable, bidirectional cross-resistance.** Grey shading highlights concentrations where cross-resistance can occur in both directions, depending on which mutant fixes. Genotypes [1 0 0] (blue) and [0 1 0] (orange) share the same dose-response curve in drug A; [0 1 0] (orange) and [0 0 1] (green) share it in drug B. Curves are slightly offset for clarity.

There can be two reasons why adaptation to drug B does not lead to cross-resistance to drug A. First, competition happens: a high-resistance mutant is likely to emerge and outcompete the low-resistance mutant before it reaches fixation (orange MSW for drug B in Figure 3a). Second, the low-resistance mutant is not adaptive at higher drug concentrations where the high-resistance mutant is adaptive (grey MSW for drug B in Figure 3a). In either case, the low-resistance mutant that would have cross-resistance to drug A is not present in the final population adapted to drug B. In contrast, adaptation to drug A will always involve a genotype that exhibits cross-resistance to drug B. Hence, this defines unidirectional, repeatable cross-resistance.

In the absence of a cost of mutations, the double mutant could sometimes fix by drift, in which case this scenario would become non-repeatable cross-resistance in the direction of drug B *→* A, but remain repeatable cross-resistance in the direction of drug A *→* B.

Figure S2 shows a scenario with higher cross-resistance to drug B following adaptation to drug A than resistance to drug B following adaptation to drug B, and Section 6.2 explains this scenario and its dependence on the drug concentration. Other explanations for unidirectional cross-resistance, following the same logic or a different one are found in Sections 6.2 and 7.

#### Unidirectional, repeatable collateral sensitivity

is easier to explain: a single mutation confers resistance to drug A and simultaneously causes collateral sensitivity to drug B. In contrast, resistance to drug B arises from a different mutation, which does not alter the MIC for drug A (Figure 2d). Hence, adaptation to drug A leads to collateral sensitivity to drug B, but not vice versa.

### 3.3 Non-repeatability relies on different resistant genotypes having different collateral effects

Several experimental studies have demonstrated that collateral effects may vary when adapting to one drug — even when experiments are conducted under identical conditions and from the same parental strain (Nichol et al., 2019; Barbosa et al., 2017). Non-repeatability requires that different resistant genotypes can fix, and at least some of these genotypes have distinct collateral effects.

#### Non-repeatable, bidirectional cross-resistance

means adaptation to drug A or B does not consistently confer cross-resistance to the other under the same conditions. This variability in the outcome can be explained by assuming that at least three loci can be mutated: two that confer resistance to drug A and two possible mutations that confer resistance to drug B, with one mutation in common, conferring resistance to both drugs (Figure 2e).

For example, in the grey MSW in Figure 3b, non-repeatability can arise when selection is strong enough, and mutations are rare enough for either resistant single mutant to fix before the other one arises. Alternatively, if mutations are more frequent or selection is not strong enough, drift, rather than selective sweeps, must ensure fixation for non-repeatability to occur. This can happen when both single mutants have similar fitness or the population size is small.

Note that the scenario described above assumes that the double mutant has a higher cost than a single mutant. Without a cost of resistance, the double mutant could drift in frequency. As the double mutant has a collateral effect, for this scenario to remain non-repeatable, the double mutant should not always be present.

#### Non-repeatable, bidirectional collateral sensitivity

can be explained in the same way. In the case where adaptation to one drug confers *no collateral effect* or collateral sensitivity to the other drug, at least four loci are needed. Two mutations confer resistance to drug A, only one of them also confers collateral sensitivity to drug B and the other does not give a collateral effect to drug B. Another two mutations do the same for drug B. These scenarios, for bi-/unidirectional non-repeatable collateral sensitivity, can be found in Figure 2g and h in the top left. However, fewer loci can explain this pattern, when adaptation to one drug confers *cross-resistance* or collateral sensitivity to the other drug (2g and h in the bottom right and discussed further in Section 6.2).

#### The degree of repeatability

, or the probability of exhibiting a particular collateral effect, can depend on various factors. First, it depends on the number of possible genotypes that confer resistance and have different collateral effects, among which the pathogen cannot transition without a decrease in growth rate (i.e., fitness peaks) and on how easily they can be reached from the parental genotype. Second, it depends on the probability of clonal interference rather than selective sweeps. Third, it is influenced by the probability of a mutation at each locus (Delaney et al., 2025). Bailey et al. (2017) found that larger population sizes correlate with higher repeatability. Finally, genetic drift and drug concentration also play a role.

## 4 Temporal dynamics of collateral effects

Earlier, we defined the collateral effect when the genotype composition has stabilised; now we define temporal dynamics as the dependence of the collateral effect in drug B on the time that the population was allowed to adapt to a constant concentration of drug A.

One example occurs when the double mutant is either the fittest or equally fit as other mutants for certain drug concentrations. This is the case, for example, in Figure 4a for drug A, when dosing at concentrations where the double mutant is the fittest. Here, the population can go through the single low-resistance mutant [0 1], which shows cross-resistance to drug B, towards the double mutant, which still shows cross-resistance to drug B (this depends on how mutational effects combine and how large both single effects are). Alternatively, the population can go through a single high-resistance mutant [1 0] that shows collateral sensitivity to drug B, and then evolve into a double mutant that shows cross-resistance to drug B. Thus, depending on when the population would be phenotyped, a different collateral effect or a mix could be observed.

**Figure 4:**
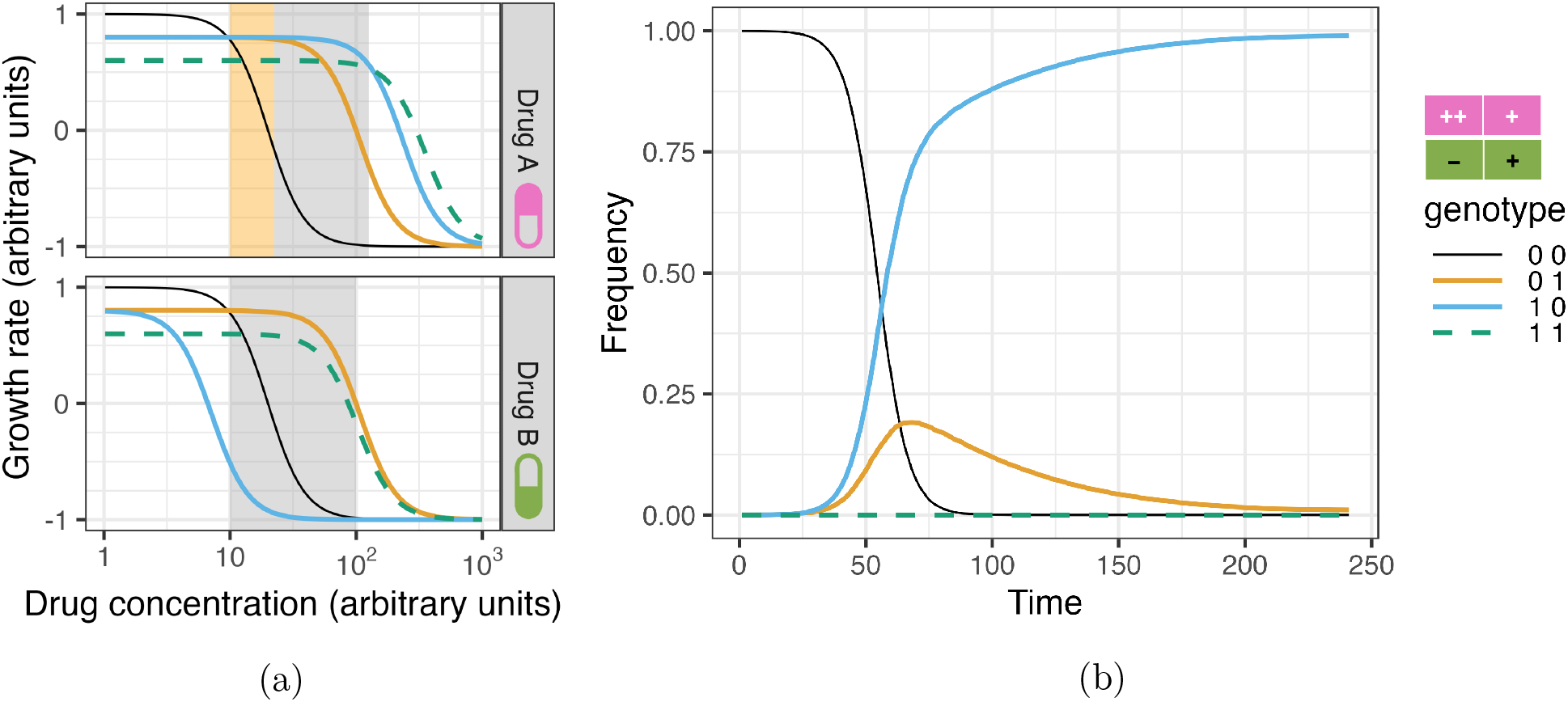
Mixed cross-resistance and collateral sensitivity and competition can lead to transient collateral effects. In the mutation-MIC shift table (top right), columns represent mutable loci; rows indicate drug-specific effects (+ means resistance, ++ means greater resistance, - means increased susceptibility compared to wildtype). **(a) Dose-response curves for two drugs with mixed collateral effects.** The grey mutant selection window denotes drug concentrations that can lead to collateral sensitivity in one direction (A *→* B) (blue, [1 0] genotype) and will lead to cross-resistance in the other direction (B *→* A) (orange, 01 genotype) (reasoning is found in Section 3.2). The orange MSW can produce collateral sensitivity, cross-resistance, or both phenotypes coexisting (A *→* B). **(b) Time series of competition dynamics between wildtype and both single resistant mutants** when dosing with 60 AU of drug A (within grey MSW) and starting from 1e6 wild type bacteria and 1e2 of both single mutants. No mutations were allowed. Intrinsic death rate was set at 0.00007 and carrying capacity at 1e9. Simulations used the code of Trubenová et al. (2022).

Another example depends on competition between single mutants. Assume a scenario where one resistance mutation confers cross-resistance, and another mutation confers collateral sensitivity to another drug (Figure 4a, time course in Figure 4b). Within the grey mutant selection window, the fittest mutant in drug A, genotype [1 0], shows collateral sensitivity to drug B. In a selection regime where competition between mutants is expected, both single mutant strains, [0 1] and [1 0], will coexist at some times. Thus, part of the population will show collateral sensitivity, and part will show cross-resistance to drug B. This would be transient until the fitter population, here corresponding to collateral sensitivity, outcompetes the less fit population (Figure 4b).

In both cases, the observed collateral effect would depend on the length of the experiment when collateral effects were quantified and on which clones were phenotyped, as the population is not always clonal.

## 5 Implications of changing selection regime or drug concentration

### 5.1 Mutation rate and selective sweeps can change directionality patterns

Unidirectional cross-resistance can be explained by the presence of both low-and high-resistance loci. Before, we assumed either a competition selection regime or a high drug concentration where the low-resistance mutant does not survive, see Figure 3a and Section 3.2. However, in a context where mutations are rare, clonal interference is reduced, thereby favouring selective sweeps. Within the mutant selection window where both single mutants outcompete the double mutant as well as the wild type, when selective sweeps occur, either of the single mutants may fix in the population.

Because these mutants have different collateral effects, the resulting cross-resistance is non-repeatable when evolving in drug B followed by exposure to drug A (B *→* A) but remains repeatable in the other direction (A *→* B) (Figure 3a). While the genetic basis is the same as in the scenario of unidirectional, repeatable cross resistance shown in Figure 3a, the change in the selection regime, for the same MSW, altered the repeatability and directionality pattern to become bidirectional: non-repeatable in one direction and repeatable in the other.

An example of the selection regime affecting the directionality of collateral sensitivity requires extending the minimal mutation-MIC shift map seen in Figure 2d; an example is found in Figure S5.

### 5.2 Increasing drug concentration can change repeatability patterns

Increasing the drug concentration can make the double mutant constitute the fittest genotype, depending on how mutational effects combine (see Figures 3a and 3b). In such a case, if the population avoids extinction, a double mutant will eventually arise and outcompete the rest, fixing in the population. This has two consequences. First, the evolutionary outcome from Section 3.3, which was non-repeatable, becomes repeatable. The double mutant shows a collateral effect to the other drug, hence this is repeatable if enough time is allowed to pass.

Second, unidirectional cross-resistance from Section 3.2 becomes bidirectional, repeatable cross-resistance. A similar case is shown in Figure S5 for collateral sensitivity.

## 6 Empirical observations

### 6.1 Validation, drug concentration and selection regimes

While many experimental evolution studies reported the resistance level (e.g. MIC, IC90) and mutations relative to wild type in their antibiotic resistant strains, the full drug response against all drugs (dose-response curve) was not measured for any or only for some intermediate genotypes (Sakenova et al., 2025; Waller et al., 2023; Barbosa et al., 2017; Imamovic et al., 2018). Other studies have systematically mutated specific genes involved in drug resistance to get a combinatorially complete set of genotypes. They then quantified the drug response of these genotypes for various concentrations of one drug (Papkou et al., 2023; Das et al., 2020; Palmer et al., 2015; King et al., 2025) or multiple drugs (Ogbunugafor et al., 2016; Mira et al., 2015; Costanzo et al., 2011) but have not related this to patterns of collateral effects during drug resistance evolution. To get a comprehensive understanding of the evolutionary dynamics driving collateral effects and their patterns, we should combine both approaches with an understanding of mutational supply to assess the selection regime.

Experimental evolution studies have looked into selection regimes by varying the concentration time series or the population size. Changing only the concentration gradient did not affect collateral effect profiles (Brepoels et al., 2022) or did so inconsistently across population sizes and bacterial species (Chauhan et al., 2025). Changing the concentration gradient and the maximal drug concentration does not give consistent results across studies: Oz et al. (2014) finds more cross-resistance and collateral sensitivity with stronger selection gradients, whereas Jahn et al. (2017) does not. Sanz-Garćıa et al. (2020) found drug-pair dependant changes in collateral effects as the drug concentration to which bacteria were allowed to adapt increased. These experimental results could be explained by measuring the dose-response curves of intermediate and final genotypes and by tracking lineages. Brepoels et al. (2022) already went in this direction by sequencing the population at multiple time points in their evolution experiment and showing a clonal interference selection regime.

### 6.2 Meta-analysis

We conducted a meta-analysis and found 6 studies where experimental evolution of at least one bacterial strain was repeated in different antibiotics, and the response of the adapted strain was measured against these same antibiotics (Barbosa et al., 2017; Li et al., 2026; Maltas and Wood, 2019; Maltas et al., 2025; Sakenova et al., 2025; Waller et al., 2023). This means that directionality and repeatability of cross-resistance and collateral sensitivity could be assessed. Additionally, two papers where repeatability could not be assessed were included Imamovic and Sommer (2013); Imamovic et al. (2018). A few things must be noted: different studies repeated evolution a different number of times, the diversity in phenotypes of adapted strains was not always assessed (sometimes only one colony was phenotyped), and different studies used different bacterial species, strains, and experimental techniques.

One observation of this meta-analysis is that cross-resistance (A *→* B) can sometimes be higher than resistance (B *→* B) (Figure 5). Several explanations are possible; we hypothesise a few that rely on having adapted only to a lower concentration of drug B when directly exposed to it, even though adaptive genotypes for higher concentrations of drug B exist.

**Figure 5:**
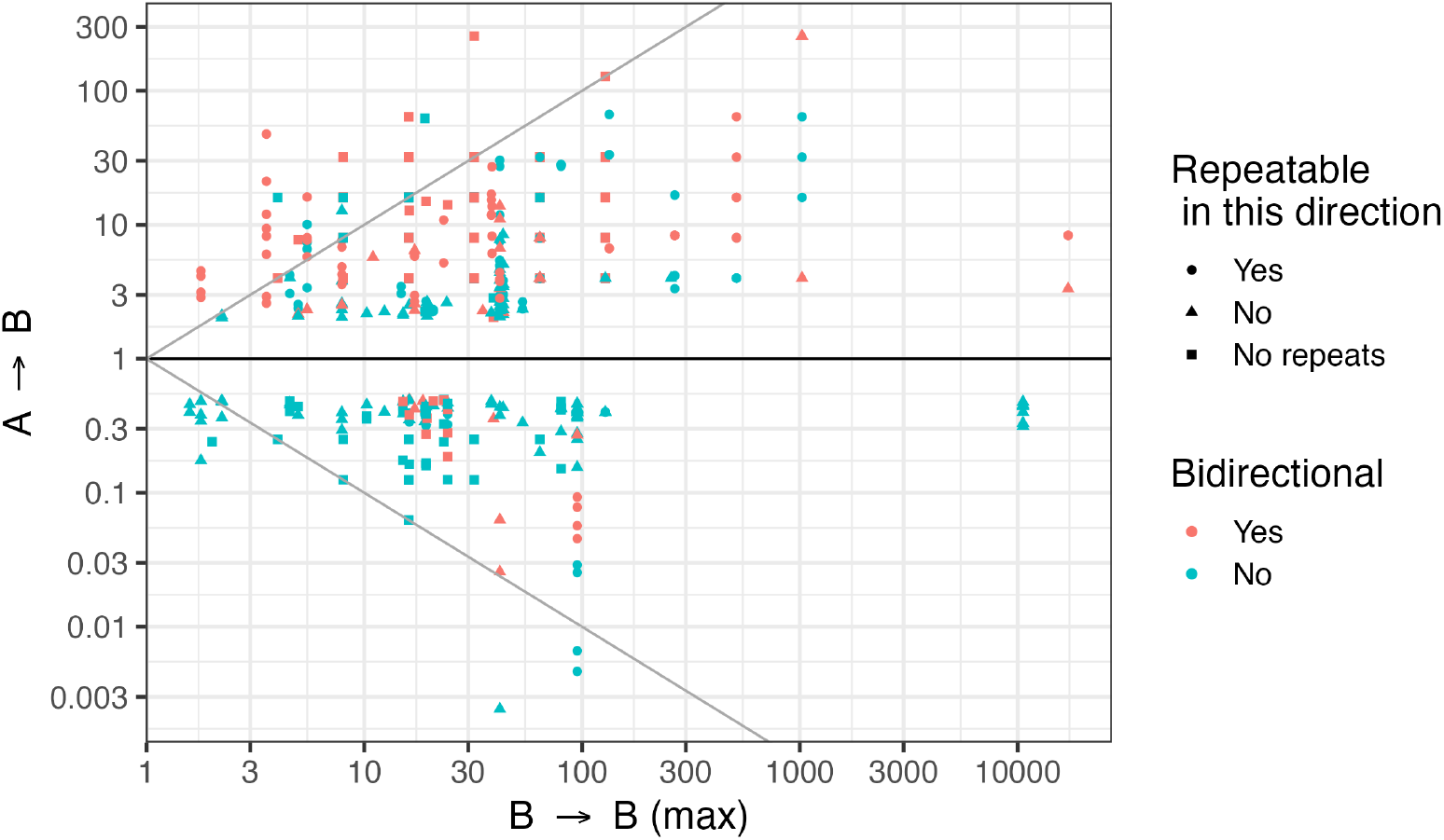
MIC or IC90 fold change to one drug when adapted to the same drug (x-axis) or another drug (y-axis). Meta-analysis of experimental evolution of different bacteria to different antibiotics. This plot includes data from Imamovic et al. (2018); Imamovic and Sommer (2013); Li et al. (2026); Maltas and Wood (2019); Maltas et al. (2025); Waller et al. (2023) as they measured the necessary MIC or IC90 values. The x-axis shows the maximal MIC or IC90 fold change to drug B achieved over all repeats of adapting the same parental strain to drug B, if repeats were done. Cases without a collateral effect are excluded. Values greater than a 2-fold change in MIC, indicate resistance, whereas values lower than 0.5 indicate collateral sensitivity (Sakenova et al., 2025). Identity lines y=x and y=-x are added to guide the eye. The data of (Barbosa et al., 2017) is found in Figure S4.

One possible explanation is that the high resistance mutation has a higher cost of resistance than the low resistance mutation (Figure S2 and Figure 2b bottom right). This phenomenon is observed experimentally (Melnyk et al., 2015). This implies that there is a MSW for drug B where the lower-resistance mutant outcompetes the high-resistance mutant. Depending on whether this low-resistance mutant shows cross-resistance to drug A or not, this scenario is bidirectional or not (the unidirectional scenario is shown in Figure S2). The second possibility is that the high-resistance mutant has a less steep dose-response curve than the low-resistance mutant, effectively creating a scenario with MSW similar to that in the increased-cost scenario described above. Both of these explanations require a clonal interference selection regime. Thirdly, there could be more low resistance mutations, or they could occur with higher probability, than high resistance mutations to drug B. Thus, the high-resistance mutant would not be frequently observed under drug concentrations where the low-resistance mutants are adaptive. This would imply that if experimental evolution were repeated more times, this high-resistance mutant would appear.

Finally, another explanation accounts for the gradually increasing drug concentration used in experimental evolution. It assumes that adaptive mutations to lower concentrations of drug B have strong negative epistasis with a particular high-resistance mutation to drug B. This means that this particular high-resistance mutation to drug B would not be observed when adapting to drug B during experimental evolution. If these adaptive mutations to lower concentrations of drug B are not adaptive to drug A, but this high-resistance mutation is, the low-resistance mutations to drug B would not be selected for when adapting to drug A, while the high-resistance mutation to drug B would. This implies higher cross-resistance than resistance to drug B.

The presence of higher cross-resistance than resistance highlights that evolution experiments should be run at clinically relevant drug concentrations. This avoids selecting for genotypes that lack cross-resistance at lower drug concentrations (unidirectional cross-resistance) or avoids selecting for lower cross-resistance than is possible (bidirectional cross-resistance). Depending on the explanation, it also advocates for repeating evolution experiments under the same conditions.

We proceeded to investigate how often both collateral sensitivity and cross-resistance were observed for different populations that were adapted to the same drug A and exposed to the same drug B (data from (Waller et al., 2023; Maltas and Wood, 2019; Maltas et al., 2025; Barbosa et al., 2017; Li et al., 2026; Sakenova et al., 2025)). This means that repeating evolution sometimes led to collateral sensitivity and sometimes to cross-resistance and thus that the collateral effect was not repeatable. This corresponds to the alternative mutation-MIC shift maps found in Figure 2, cells e, g and h bottom right, for non-repeatable uni-/bidirectional collateral sensitivity and bidirectional cross-resistance. We analysed our data independent of directionality, i.e. looking at all collateral effects to drug B found when adapted to drug A (the drug pair here is A-B). Collateral sensitivity was more often non-repeatable than repeatable for all studies except Li et al. (2026). When it was non-repeatable, only one study (Barbosa et al., 2017) found more drug pairs, 12, where repeated evolution sometimes showed cross-resistance and sometimes collateral sensitivity, and only 4 drug pairs where repeated evolution sometimes showed no collateral effect and sometimes collateral sensitivity but never cross-resistance. Three other studies (Maltas and Wood, 2019; Maltas et al., 2025; Waller et al., 2023; Sakenova et al., 2025) found that no collateral effect, not cross-resistance, was most often observed when collateral sensitivity was non-repeatable. For Li et al. (2026), one drug pair showed collateral sensitivity and cross-resistance, another showed collateral sensitivity and no collateral effect. On the other hand, non-repeatable cross-resistance, most often showed no collateral effect, not collateral sensitivity for all studies. In conclusion, non-repeatable collateral sensitivity can mean that sometimes cross-resistance is observed but often it is not and no collateral effect is observed.

## 7 Conclusion and discussion

In this study, we proposed minimal examples that use the fewest number of loci needed to describe combinations of repeatable and non-repeatable, uni-or bidirectional patterns of cross-resistance and collateral sensitivity, as shown in Figure 2. We reasoned how these patterns can change with drug concentration and selection regime. Furthermore, we demonstrated that dose-response curves for a combinatorially complete set of genotypes, measured across different single-drug environments, should reveal how specific drug concentrations and selection regimes determine the emergence of various collateral effects of drug resistance and their patterns: repeatability, directionality, and temporal dynamics.

This study omits complex epistatic interactions (though see Section S1.2). For example, epistasis could provide another explanation for unidirectional cross-resistance if we assume that multiple mutations are required for resistance to drug A, conferring resistance only when all are present (B. Vicoso, personal communication, May 13, 2025). A subset of these mutations gives resistance to drug B. Hence, if a wild type population adapts to drug A, it is also cross-resistant to drug B, but not vice versa. This is the only explanation where unidirectional cross-resistance is independent of both drug concentration and selection regime. Another limitation is the lack of a direct link between the inferred genetic architecture and the underlying molecular mechanisms of resistance and collateral effects. Furthermore, we did not consider a changing environment. For example, trade-offs between periods of presence and absence of drugs could define the fittest genotype. Another example is the particular drug concentration time course used in experimental evolution (see Section 6.2). Future directions thus include understanding how mutations combine to affect the dose-response curve (e.g. Barbosa et al. (2019) looked at epistatic interactions between mutations on MIC). The effect of a changing drug environment can be assessed experimentally (Maltas et al., 2025) but this also renders itself to simulation studies (Nichol et al., 2015).

The purpose of this study was to find a first set of examples of genotype-phenotype maps and their dose-response curves and the logical reasoning of the effect of the selection regime and drug concentration on collateral effects and their patterns of repeatability, directionality and temporal dynamics. We hope that this framework will be useful for understanding the patterns of repeatability and directionality of collateral effects for both theoretical modellers and experimental biologists studying the evolution of drug resistance, not only in bacteria, but across taxonomic boundaries.

## Acknowledgements

We acknowledge and appreciate funding from the Swiss National Science Foundation. We would like to thank Mario Bolli, Aryahi Kumar, Beatriz Vicoso, Chris Witzany and Fernando Rossine for their helpful discussions and/or suggestions that helped us better understand and formulate the described concepts. Furthermore, we thank Hinrich Schulenburg, Aryahi Kumar, Viktor Kovalov, Basil Vogelsanger, and Julian Zaugg for providing feedback on the manuscript.

## 8 Supplementary material

The code used to create the illustrations is found at the following link: https://github.com/Mayiiiiii/CollateralEffectsDrugResistance.

### S1.1 Minimal examples of special cases

There are several special, but important cases that were not discussed in the main text due to space limitations.

#### Bidirectional, repeatable in one direction, non-repeatable in the other

cross-resistance can be explained by assuming that there are two possible loci that can be mutated. Both mutated loci confer resistance to drug A, but only one confers resistance to drug B. This corresponds to the dose-response curves in Figure 3b without the third locus. Following the same reasoning and assumptions as Section 3.3, for an intermediate drug concentration range, adaptation to drug A can sometimes but not always lead to cross-resistance to drug B, and adaptation to drug B will always lead to cross-resistance to drug A. A similar case is discussed in Section 5.

#### A special case of competition that leads to unidirectional cross-resistance

(Section 3.2) occurs when the low-resistance mutant grows so slowly at a given concentration that it takes a long time to become established. During this extended period, there is a high probability that a high-resistance mutant will emerge from the remaining wild type sub-population and outcompete the low-resistance mutant. This scenario is only likely if the wild type subpopulation has a very slow net death rate and some growth. If the high-resistance mutant appears first, it can quickly become fixed. If the low-resistance mutant appears first, the slow population dynamics mean that the high-resistance mutant is still likely to arise and then outcompete the low-resistance mutant.

### S1.2 Assumptions on combining mutational effects and alternative explanations for unidirectional cross-resistance

#### Violating the assumptions in **Section 2** can also give the patterns we describe

Additive effects of multiple mutations on cost and MIC shift, as well as constant steepness (*κ*) between mutants, are not the only conditions that allow for the reasoning we used in Section 3 to describe the collateral effect patterns.

For example, **unidirectional cross-resistance** (Section 3.2), for a drug B where adaptation does not lead to cross-resistance to drug A, can be explained by:

1. two resistance mutations to drug B. One with and one without cross-resistance to drug A.
2. the drug B concentration is within a mutant selection window (MSW) where the single mutant without cross-resistance outcompetes the other single mutant, or the other single mutant is not adaptive in this drug concentration.
3. Regarding the double mutant, it also requires one of two things:

(a) If the double mutant shows cross-resistance, then the double mutant should be less fit than the single mutant without cross-resistance for the relevant MSW.

i. This is the case when the cost of resistance increases when combining these mutations (Figure 3a). However, this cost increase need not be additive.
ii. Another option is a less steep dose-response curve for the double mutant (an example is found in Figure S3: the same reasoning from Section 3.2 can be applied).
iii. The double mutant is not similarly fit or more fit than the single-resistant mutant without cross-resistance in this MSW.

(b) Alternatively, it requires that the combination of these two mutations does not lead to cross-resistance through some epistatic effect.

A second explanation is that you have a linear pathway 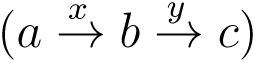 and drug A acts on the first reaction *x* and drug B acts on the second reaction *y*. If we assume that resistance to drug A emerges by bypassing these normal reactions in the following way 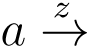 and that resistance to drug B emerges by bypassing only the second reaction 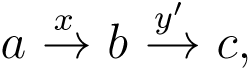, then resistance to drug A implies cross-resistance to drug B but not vice versa.

## S1.3 Supplementary figures

**Figure S1:**
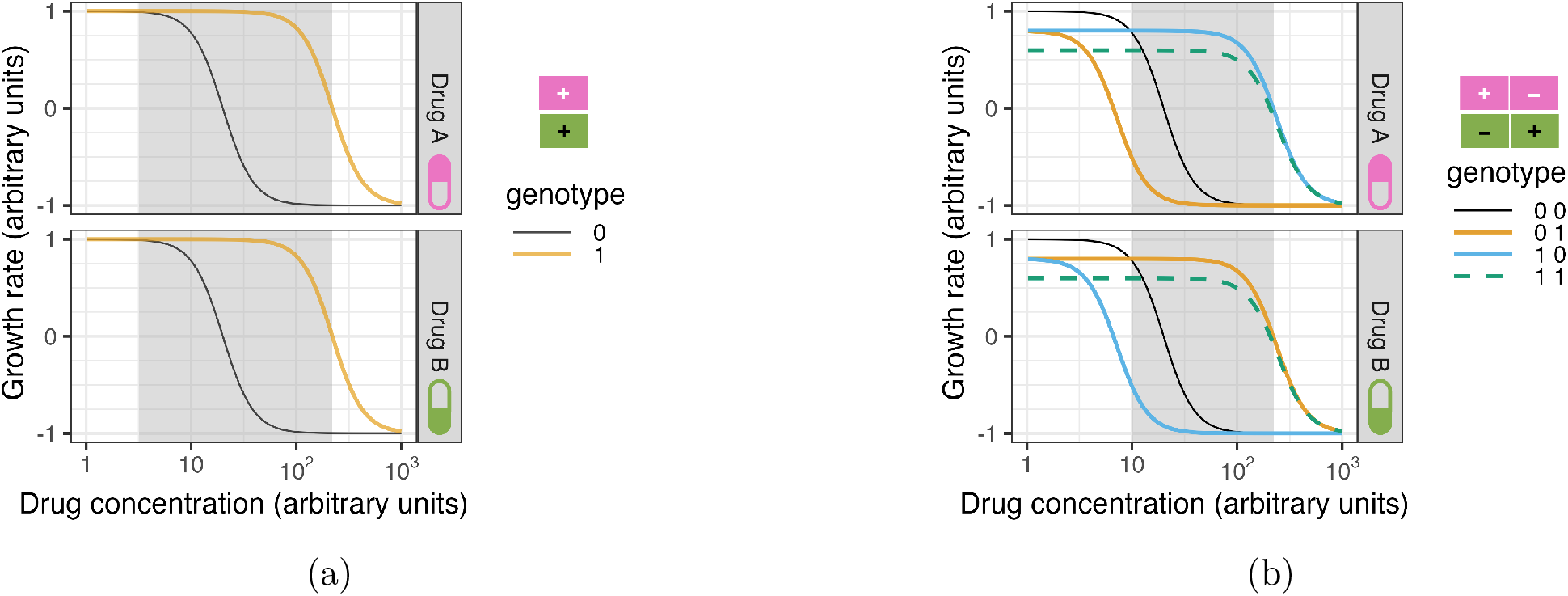
Dose-response curve examples for two drug environments with corresponding mutation-MIC shifts that can lead to different collateral effect patterns. Mutation-MIC shift: the mutation effect at this locus (column) in drug A (pink top row) and drug B (green bottom row). + means resistance compared to the wild type. - means increased susceptibility compared to wild type. (a) **Bidirectional, repeatable cross-resistance.** The grey mutant selection window denotes drug concentrations that will lead to cross-resistance if the population is adapted to either drug (orange, [1] genotype). (b) **Bidirectional, repeatable collateral sensitivity.** The grey mutant selection window denotes drug concentrations that will lead to collateral sensitivity if the wild type population adapted to either drug A (blue, [1 0] genotype) or drug B (orange, [0 1] genotype).

**Figure S2:**
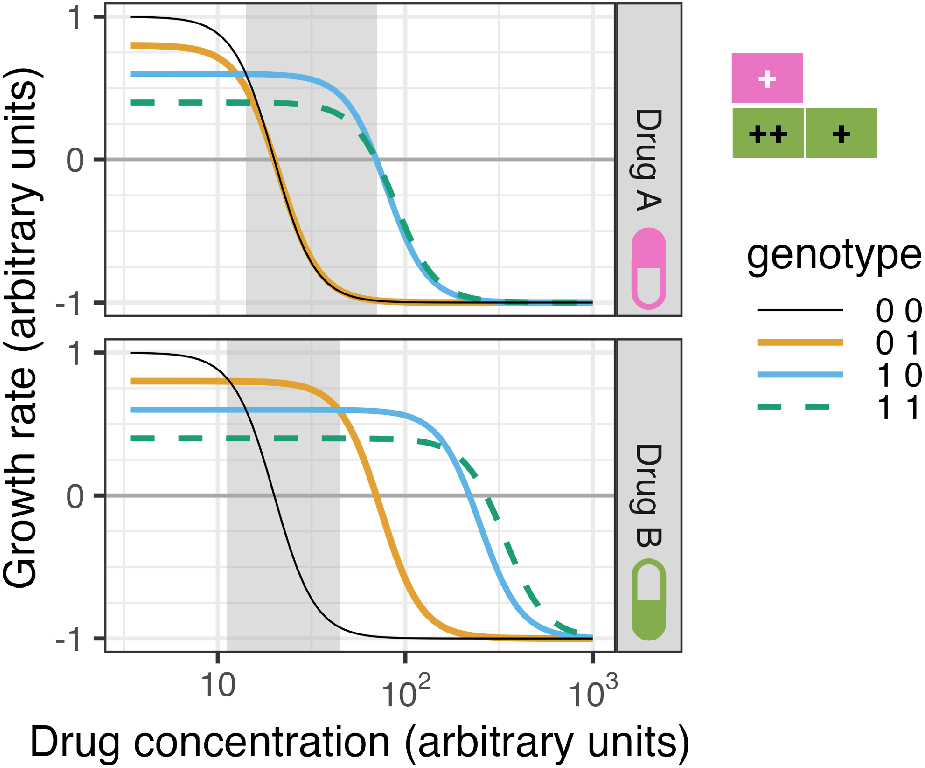
Unidirectional repeatable cross-resistance can arise when a low-resistance mutant has a higher growth rate than a high-resistance mutant at intermediate drug concentrations. Grey shading highlights the MSW where cross-resistance to drug B occurs only when adapted to drug A, and there is no cross-resistance to drug A when adapted to drug B in a clonal interference selection regime.

**Figure S3:**
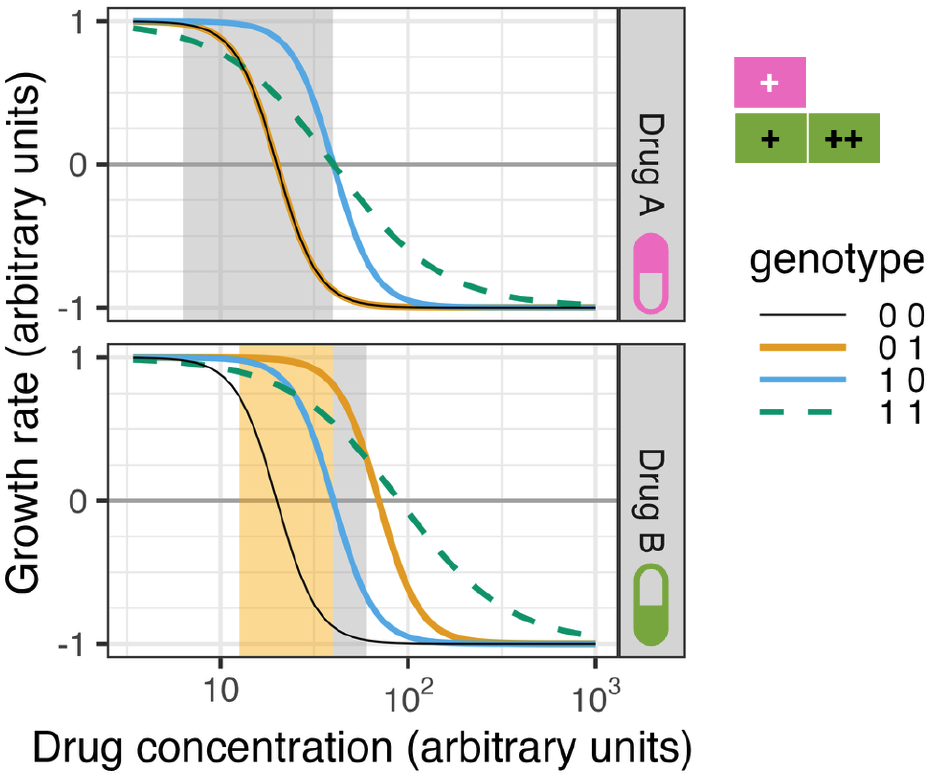
Unidirectional, repeatable cross-resistance. Dose-response curve example for two drug environments with corresponding mutation-MIC shift. Mutation-MIC shift: the mutation effect at this locus (column) in drug A (pink top row) and drug B (green bottom row). + means resistance compared to the wild type. ++ refers to a larger increase in MIC than +. The grey mutant selection windows denote drug concentrations that will lead to cross-resistance only if the population adapted to drug A first (blue, [1 0] genotype), but not if it adapted to drug B first (orange, [0 1] genotype). The orange mutant selection window for drug B is the concentration range where competition between mutant strains is necessary to always fix the genotype without a collateral effect in drug A (orange, [0 1] genotype). *κ* is 4 for all genotypes but 1.5 for the double mutant. The first locus shifts the MIC in both drugs by 20, the second locus shifts the MIC in drug B only by 50. There is no cost of resistance.

**Figure S4:**
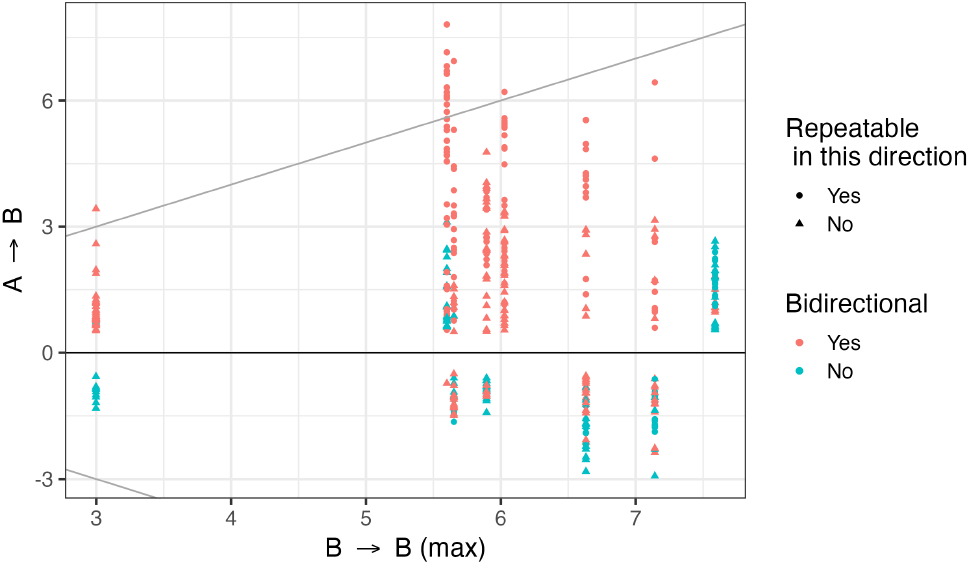
An overview of the effect size of the collateral effect to drug B when evolved in drug A compared to resistance to drug B when evolved in drug B for Barbosa et al. (2017). Cases without a collateral effect are excluded. Values greater than 0.5 indicate resistance, whereas values lower than-0.5 indicate collateral sensitivity. The x-axis shows the maximal area under the curve (AUC) difference to drug B achieved over all repeats of adapting the same parental strain to drug B, if repeats were done. Identity lines y=x and y=-x are added to guide the eye. One set of vertically aligned points represents one drug B, hence the number of such sets is the number of drugs tested.

**Figure S5:**
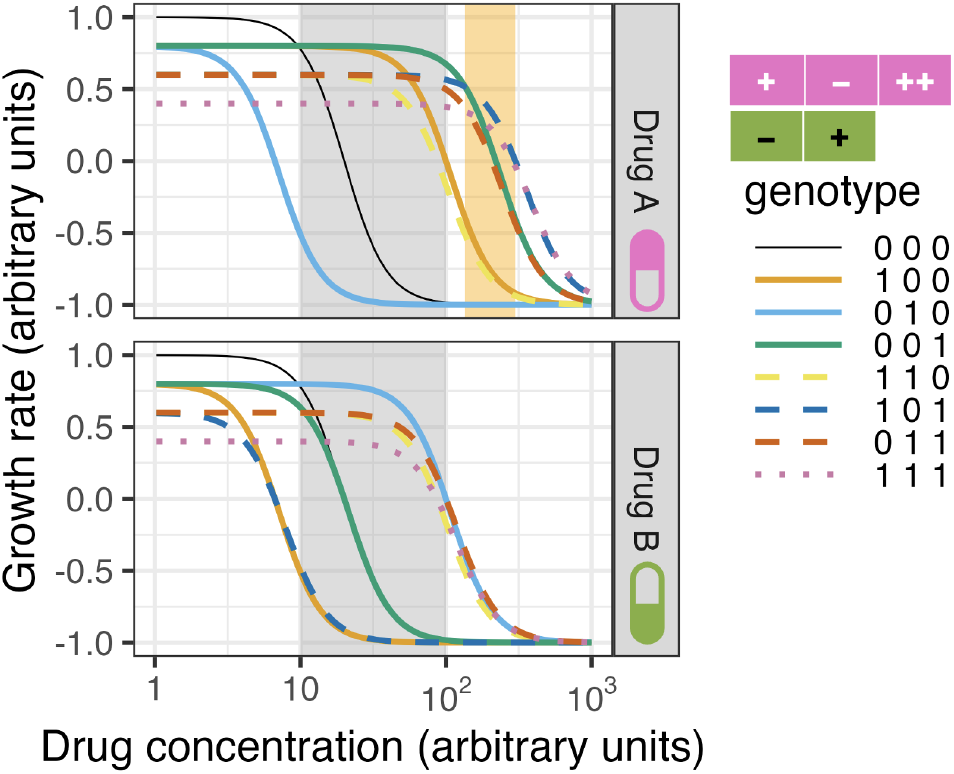
Dose-response curve examples for two drug environments with corresponding mutation-MIC shifts that can lead to **different directionality of collateral sensitivity depending on the selection regime and drug concentration**. Mutation-MIC shift: the mutation effect at this locus (column) in drug A (pink top row) and drug B (green bottom row). + means resistance compared to the wild type. ++ refers to a larger increase in MIC than +. - means increased susceptibility compared to wild type. When adapting to drug B in the grey MSW, collateral sensitivity to drug A would always be observed, genotype [0 1 0] (*B → A*). When adapting to drug A on the other hand, the selection regime and drug concentration matters. In the grey MSW, clonal interference would cause the [0 0 1] genotype to prevail and thus no collateral sensitivity to drug B would be observed. This would imply unidirectional collateral sensitivity. On the other hand, in the grey MSW, selective sweeps would mean that sometimes the [1 0 0] genotype and sometimes the [0 0 1] genotype would fixate. Depending on the exact drug concentration, certain genotypes with more mutations would be selected for after the [1 0 0] genotype sweeps. In any case, this would imply non-repeatable collateral sensitivity (*A → B*) and repeatable collateral sensitivity (*B → A*). Finally, for the orange MSW, the [1 0 1] genotype would be selected for. This genotype shows collateral sensitivity to drug B, and hence this would imply repeatable, bidirectional collateral sensitivity.

## S1.4 Supplementary tables

**Table S1:**
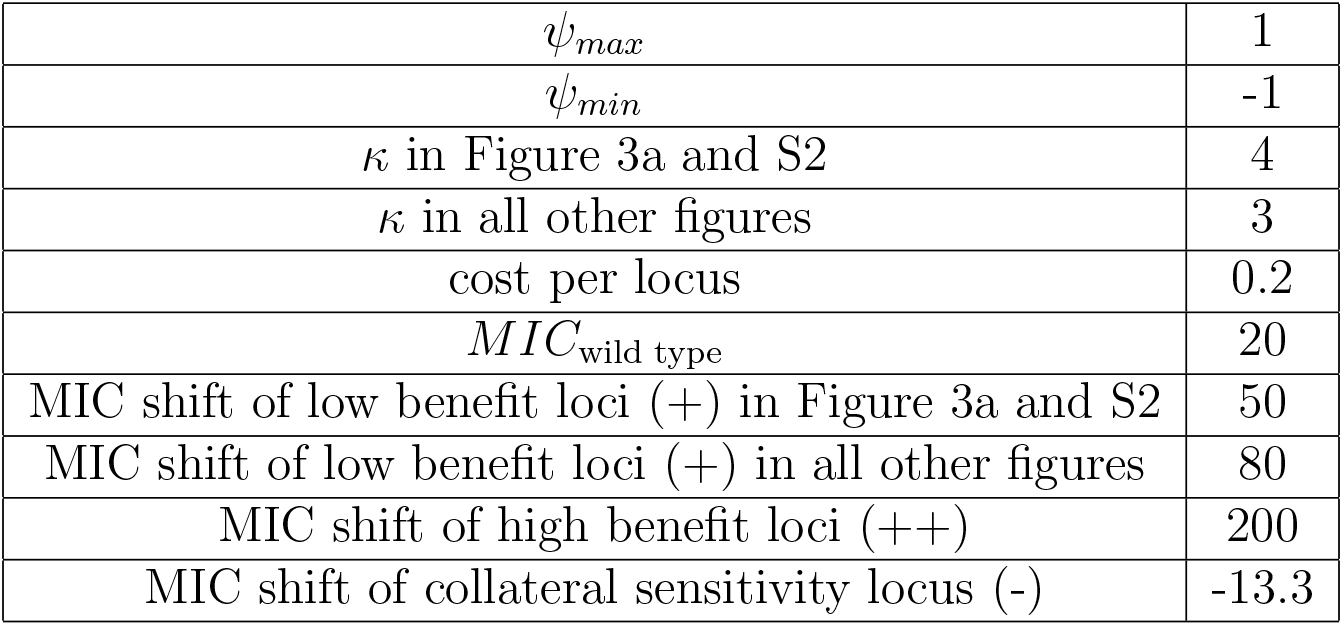
Parameters of illustrated dose-response curves. These parameters of Equation 1, used in all figures unless specified otherwise. These parameters are arbitrary and used to allow for clear illustrations of the ideas presented in this manuscript.

|  |  |
|---|---|
| ψ<sub>max</sub> | 1 |
| ψ<sub>min</sub> | -1 |
| κ in Figure 3a and S2 | 4 |
| κ in all other figures | 3 |
| cost per locus | 0.2 |
| *MIC*<sub>wild type</sub> | 20 |
| MIC shift of low benefit loci (+) in Figure 3a and S2 | 50 |
| MIC shift of low benefit loci (+) in all other figures | 80 |
| MIC shift of high benefit loci (++) | 200 |
| MIC shift of collateral sensitivity locus (-) | -13.3 |

## Notes

https://github.com/Mayiiiiii/CollateralEffectsDrugResistance

### Competing Interest Statement

The authors have declared no competing interest.

### Summary of Updates

New section on experimental observations, minor updates throughout, updated figure 2, merged boxed into text, added some supplementary figures.

